# A randomized controlled trial of the effectiveness of a community-based rabies vaccination strategy

**DOI:** 10.1101/2024.10.28.620430

**Authors:** Felix Lankester, Ahmed Lugelo, Joel Changalucha, Danni Anderson, Christian Tetteh Duamor, Anna Czupryna, Kennedy Lushasi, Elaine Ferguson, Emmanuel S. Swai, Hezron Nonga, Maganga Sambo, Sarah Cleaveland, Sally Wyke, Paul C. D. Johnson, Katie Hampson

## Abstract

**Background:** Approximately 60,000 people die from dog-mediated rabies annually. Low and heterogeneous coverage reduces the effectiveness of dog vaccination campaigns that can eliminate rabies. Campaigns typically involve teams travelling annually to villages to deliver cold chain stored vaccines from centralized headquarters. Thermotolerant vaccines enable novel decentralized delivery of locally-stored vaccines by communities throughout the year. We compared the effectiveness of annual team-based versus continuous community-based dog vaccination strategies.

**Methods:** We conducted a cluster randomized controlled trial across Mara region, Tanzania. Trial clusters were administrative wards (112, on average four villages each). For the team-based arm vaccinators hosted annual static-point clinics, whilst for the community-based arm, a ward-based animal health officer with a village community leader managed vaccinations using vaccines stored within the ward. We measured vaccination coverage, the primary outcome, twice annually per cluster (month 1 and 11) through household surveys over three years (November 2020 to October 2023) and examined spatial and temporal coverage variations as secondary outcomes.

**Findings:** Community-based delivery achieved significantly higher coverage (49-62%) than team-based delivery (22-46%), and consistently exceeded the critical threshold for herd immunity (40%), Odds ratio (OR): 1.48-3.49. The lower less uniform coverage achieved through team-based delivery had a higher monthly probability of falling below the critical threshold (0.6, 95% CI: 0.38-0.81) vs 0.18 (95% CI: 0.04-0.40). Greater declines in coverage over the year were recorded in the team-based arm compared to the community-based

**Conclusion:** Community-based mass dog vaccination achieves higher more consistent coverage than team-based delivery across settings typical of many sub-Saharan African countries. This approach could play an important role in national rabies elimination programmes aiming to end human rabies deaths by 2030 as part of the global ‘zero by 30’ strategy.

**Funding:** Department of Health and Human Services of the National Institutes of Health (R01AI141712), Wellcome Trust (207569/Z/17/Z, 224520/Z/21/Z) and MSD Animal Health. The content is solely the responsibility of the authors and does not necessarily represent the official views of the National Institutes of Health.

## Introduction

Dog-mediated human rabies kills approximately 59,000 people annually, mostly children, with millions more saved only by costly post-exposure prophylaxis (PEP)^1,2^. The vast majority (>99%) of human rabies fatalities occur in Africa and Asia, where access to PEP is limited^1,3^. The World Health Organization (WHO), the Food & Agricultural Organization (FAO) and the World Organization for Animal Health (OIE) have recognized human rabies as a global health priority and have committed to end human rabies deaths due to dog-mediated rabies by 2030^4^. Mass dog vaccination (MDV) is an efficient approach to eliminating human rabies by inoculating the reservoir host responsible for maintaining disease and causing more than 99% of human infections. However, implementing MDV across the rural landscapes is logistically challenging and expensive^5,6^. Moreover, there has only been limited empirical evidence to demonstrate the cost-effectiveness of MDV in achieving public health outcomes. As a result many countries spend substantial resources on provision of PEP with only limited investment in MDV, and human deaths continue to occur at an unacceptably high rate^1,7^.

The standard MDV delivery method across Africa is a pulsed centralized team-based delivery strategy. Centered upon hub locations where reliable power supplies allow bulk vaccine storage under cold-chain conditions (4°C), teams of vaccinators drive out to remote target villages and set up temporary static-point MDV clinics in a convenient position in the village that typically last for one day^5,8^. To eliminate rabies, these once-per-year campaigns must vaccinate at least 70% of each community’s dog population to maintain coverage above 20 – 45%, the critical threshold for herd immunity^9^. Otherwise turnover in the dog population leads to drops in coverage that allow sustained rabies transmission. Achieving this coverage consistently across remote landscapes with team-based delivery is challenging^10,11^. For example, community engagement is vital for success, yet this critical factor is influenced by agricultural cycles, school holidays and even inclement weather^5,12,13^. Accommodating these factors within the scheduling of centralized team-based strategies is logistically demanding, and as a result gaps in coverage easily arise. This is problematic as even small gaps can hinder progress to elimination^11^. Novel, cost-effective MDV delivery strategies that maintain consistently high coverage at the scale required for regional elimination are urgently needed.

Decentralized community-based strategies are a promising way of improving the delivery interventions and have been used in the control of neglected tropical diseases such as onchocerciasis^14^. A community-based model for rabies control is hypothesized to improve coverage consistency and reduce delivery costs^10,15,16^. The barrier to implementing and testing community-based delivery has been the inability to store rabies vaccines under cold-chain conditions in rural communities^17^. The availability of a thermotolerant rabies vaccine that can be stored without loss of potency for extended periods at temperatures exceeding cold-chain conditions allows the investigation of community-based delivery options. Indeed, thermotolerant formulations have been used to deliver vaccines in hard-to-reach communities in Africa and Asia, including childhood vaccines for meningitis^17^ and hepatitis^18–22^ and Newcastle disease vaccines for chickens^23^. Moreover, thermotolerant formulations that allowed local communities to deliver vaccination campaigns played a pivotal role in the eradication of both smallpox and rinderpest^24,25^. By enabling community-based delivery, a thermotolerant rabies vaccine could have the same transformative effect accelerating progress towards eliminating dog-mediated rabies.

A widely used canine rabies vaccine (Nobivac™ Rabies) can be stored remotely in locally made cooling devices while maintaining protective immunogenicity^26–28^ and that trained community operators can effectively administer vaccines^16,29^. These findings enable investigation of the effectiveness of community-based dog vaccination using continuous decentralized strategies. Here, we report results from a large-scale randomized controlled trial (RCT), that aimed to evaluate the effectiveness of a decentralized community-based delivery strategy against the standard, annual team-based delivery.

## Methodology

A stratified parallel cluster RCT was conducted in six districts within the Mara region, northern Tanzania (Figure 1). Each district consists of eight to 30 administrative wards each comprising around three to four villages. The RCT covered 112 wards of which 90 (80%) were classified as rural and 22 (20%) as urban^30^. The human population in the study area at the trial start (2020) was 1,096,136 projected from the 2012 national census^30^, while the dog population was estimated to be 183,000 from the human-to-dog ratio of 6:1^31^.

**Figure 1.**
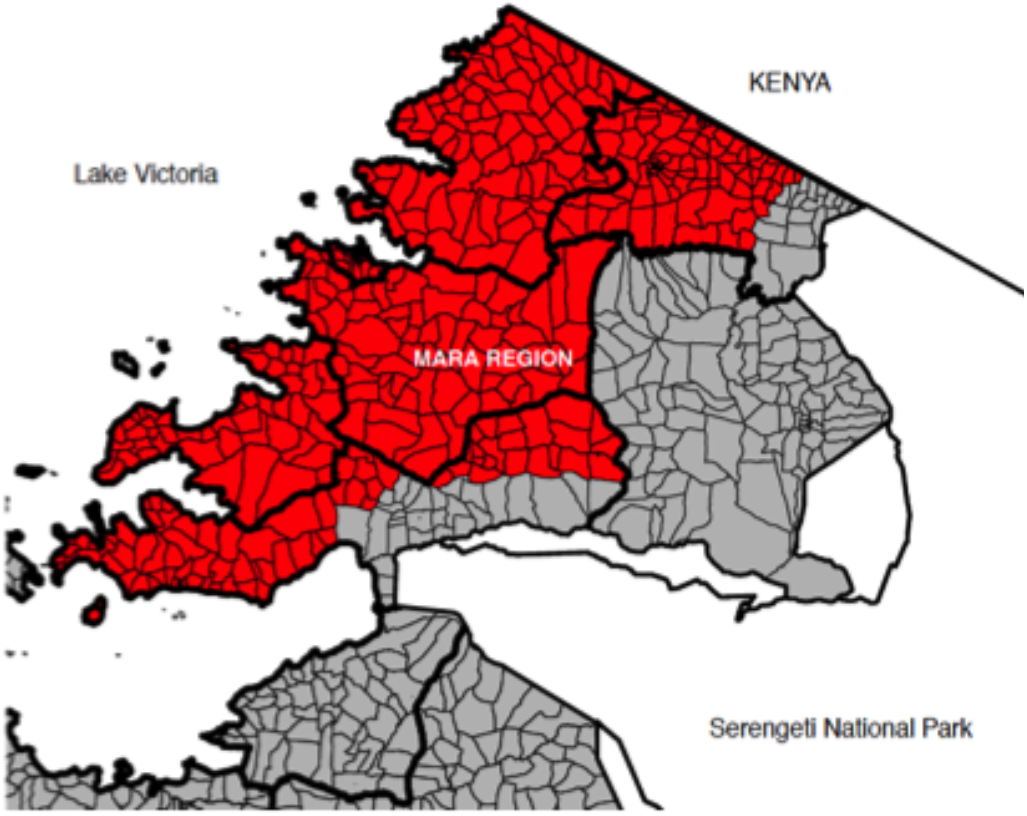
Study area in Tanzania. Districts and wards are shown with thick and thin black lines, respectively. The RCT took place in target districts colored red. Grey area indicates wards where on-going MDV had recently targeted

We tested two dog vaccination delivery strategies over three annual cycles (November 2020 – October 2023): 1) standard centralized team-based delivery, with vaccinators traveling from a central location to each target community (Arm 1) and 2) novel decentralized community-based delivery, implemented by local animal health officers and community leaders (Arm 2) The trial was clustered at the administrative ward level with each cluster being randomly selected into one of the two arms. We employed a ‘fried-egg’ approach^32^ to prevent contamination (spill-over) of intervention effects between the trial arms. The whole cluster received the allocated treatment but coverage was measured only in the inner area of the cluster (‘egg-yolk’) i.e., ward’s central-most village (study village). Consequently, the trial was powered at the study village level. Randomisation was implemented using a custom R script ^33^ written by the trial statistician.

### Inclusion/exclusion criteria

Only wards that had not previously been targeted by MDV campaigns (n = 178) were eligible for inclusion (n = 112; Figure 1, red). Wards with ongoing dog vaccination campaigns were excluded (Figure 1, grey).

### Interventions being tested

#### Team-based delivery

This strategy involved the District Veterinary Officer (DVO) forming a vaccination team consisting of Senior Livestock Field Officers (SLFOs) from a district headquarter and a ward-based Livestock Field Officer (LFO) and a village-based assistant operating in all Arm 1 villages assigned to them within their district.

The vaccination schedule included hosting a single static-point clinic in a central point of every village to begin within the first month (November) of the three annual cycles. Prior to each static-point clinic, village members were sensitized about the forthcoming MDV through their community leaders.

On each vaccination day, the SLFO travelled from the district headquarter to the predetermined village using a motorbike carrying vaccination equipment including sufficient doses of vaccines previously stored in the district refrigerator. The SLFO was responsible for the registration and inoculation of dogs, whilst the ward-based LFO completed the vaccination certificates for dog owners, and the village assistant recorded vaccination information in a register, including the dogs name, sex, age, and color and owner’s name, all of which were entered into a digital data collection survey tool by the SLFO. Following registration, all dogs received a sub-cutaneous 1-mL injection of Nobivac® Rabies vaccine.

Each village-level clinic operated for a full day. The following day, the SLFO moved to the next village, continuing until all villages in a ward were vaccinated. Then the SLFO targeted their next ward, repeating until all wards were completed. Each ward was revisited annually in the first month of the next cycle, inviting all dogs for vaccination regardless of previous status.

#### Community-based delivery

batches of vaccine as well as other vaccination equipment were managed within each ward by a ward-based animal health extension officer, thereafter called the Rabies Coordinator (RC). RCs received training on how to organise and host MDV activities. Prior to the onset of activities, the RC travelled by motorbike to the District Veterinary Office to collect vaccination equipment and a batch of Nobivac® Rabies vaccines, which had been stored under cold-chain conditions. The number of vaccines collected were estimated to be sufficient for three months of vaccination activities based upon the local dog population size, which was in turn estimated by a village-based assistant known as One Health Champions (OHC) in consultation with community leaders. RCs stored these vaccines in a locally made low-tech cooling device^27^. Prior to each village-based event, advertising was implemented by the RC and the OHC, chosen by the community and the RC from the village leadership. As with Team-based delivery, this involved posters being strategically placed in key public areas such as schools and market places. On the day before the campaign, the OHC traversed the village, using a loudspeaker to announce that the RC would vaccinate dogs the next day and encouraging people to bring their dogs. On the vaccination day, the RC travelled by motorbike to the target village carrying sufficient doses, consumables, and equipment (syringes, needles, etc.) for the day’s activities. The vaccination activities lasted for a day. The following day, the RC targeted the next village in the ward, collaborating with the village-based OHC and employing the same approach until all villages within the ward had been targeted.

As with Team-based delivery, the vaccination team consisted of: the RC responsible for the inoculation and registration of dogs, the OHC in charge of certification, and a local helper tasked with recording vaccination data in the register. Dogs brought to the clinic were registered and vaccinated in the same manner as in the Team-based arm.

A key difference between the delivery methods was that because of local storage of vaccines, vaccination takes place throughout the year under community-based delivery. To this end, in the weeks following completion of the first round, each OHC determined (through their local knowledge and discussion with community members) which dogs in their village might have been missed and whether litters had been born since. This information was communicated to the RC, with the intention that these dogs would receive vaccination during the RC’s routine animal health duties. At month three, six and nine of each year further vaccination activities were carried out by the RC in their respective villages, either as static point clinics hosted at sub-village level or house-to-house visits. The method used was chosen by the RC and contingent on the number of dogs identified that needed vaccinating. In year two and year three the procedure was repeated, with the RC instructed to vaccinate all dogs each year, irrespective of whether they had been vaccinated previously.

### Vaccination Coverage Assessment

Household surveys were conducted twice annually (Timepoints 1 and 2) in study villages to evaluate vaccination coverage in both arms from Years 1 to 3. The six household survey visits are hereafter abbreviated to Y1V1, Y1V2, …, Y3V2. The first survey, which allowed assessment of the immediate post-vaccination coverage, was scheduled as soon as possible after completion of the static-point clinic in each ward. The second survey allowed evaluation of how coverage had changed during the year and similarly was scheduled to take place between months ten and twelve. The survey team was composed of an interviewer and local guide (a resident appointed by a community leader) who provided guidance on the location of and introductions to households. Study villages were surveyed in both trial arms. In each study village three sub-villages were randomly selected before each survey (using an R script) and a target of ten dog-owning households per sub-village were surveyed as follows: a household located on the periphery of the chosen sub-village was selected as the index household from which the survey would commence. From this household the survey team walked towards the sub-village centre and to the opposite side of the sub-village. If a sub-village was targeted more than once within the three-year trial, a different index household was chosen from previous surveys. Every fifth household was chosen and if no adult was present or no dogs were owned the household was skipped and the next household was targeted. Interviews continued until ten households were surveyed in each sub-village, before continuing to the remaining sub-villages, repeating the household selection and survey procedure until the target sub-villages (n=3) and households (n=30) had been completed. If 10 dog-owning households were not found in a targeted sub-village, where feasible, additional sub-villages were added to reach the target of 30 dog-owning households.

Prior to administration of the household survey, permission was sought from the household head or, in their absence, another adult household member (> 18 years). Interviews were carried out in Swahili and, if needed, translated into the local vernacular language by the local guide. Interviewers explained the study background to each respondent and obtained written informed consent for the questionnaire. The questionnaire captured the total number of dogs living in the household, and their vaccination status. If dogs had been vaccinated, a certificate was requested, and the registration number recorded. If the owner stated that the dog had been vaccinated but no certificate could be produced, this was recorded. If there were unvaccinated dogs at the household, reasons for not vaccinating were recorded. In addition, opinions about the quality, accessibility and availability of the dog vaccination services provided were collected.

### Training

Prior to commencement of MDV activities, the research team led a series of district-level training programs aimed to familiarize the DVOs and equip implementers in both arms of the trial with the skills to perform their duties. Parallel training activities were conducted, with one series focusing on the implementers of team-based delivery and the other on implementers of community-based delivery.

In the team-based delivery training, vaccinators were trained how to advertise the MDV in advance of each village campaign, manage cold-chain stored vaccines, set up static-point clinics, receive and register dogs and owners, administer vaccinations and record vaccination data. In the community-based delivery training, the RCs were additionally trained how to collect batches of vaccines from the central storage at the respective District Veterinary Office, store vaccines within their respective wards using the low-tech cooling system, and organize and manage their vaccination campaigns in each of the villages of their wards. Additionally, the village-based OHCs were instructed on how to estimate the dog population in their respective village, identify unvaccinated dogs, and sensitise the community.

### Statistical Analysis

Analyses were performed using a generalized linear mixed-effects regression model (GLMM), which fitted using maximum likelihood, except for Secondary Analysis 4 where the GLMM was fitted using Markov Chain Monte Carlo (MCMC). Logit-normal random effects were fitted to account for spatial autocorrelation in coverage at four levels: districts, wards, subvillages, and households. When fitting the district-level random effect, three districts were split into two, giving nine levels: Bunda (Bunda Town Council, Bunda District Council); Musoma (Musoma, Musoma Municipal); and Tarime (Tarime Town Council, Tarime District Council). Two types of random effect were fitted at each level except household: location-specific random effects that allowed variation among locations that was consistent over time and time-point-specific random effects to allow variation in coverage that was not consistent over time. Because only approximately a third of households were resampled between survey visits, only a time-point-specific random effect was fitted for household. Intervention arm (Team-based and Community-based), survey visit (V1 and V2) and year (Y1-Y3) were modelled as categorical fixed effects. All three two-way interactions and a three-way interaction between year, survey visit, and trial arm were fitted. The effect of these interactions was to allow coverage to differ between the six time-points (year × visit), and the intervention effect to differ between the six time-points (arm × year, arm × visit, arm × year × visit). To protect the primary analysis from inflation of the type I error rate due to multiple testing, no variable selection was performed on the primary analysis model; specifically, all pre-specified interaction effects and main effects were retained regardless of significance. For secondary analyses 2, 5 and 6, models were simplified by backwards elimination of non-significant terms (see Table S1 for details of all analyses and models).

Null hypotheses were tested using likelihood ratio tests and rejected at the 5% significance level if P < Intervention effects were estimated from the primary analysis GLMM as community:team odds ratios. Coverage was also estimated from the GLMM. Estimates ± 95% confidence limits were calculated as logit^-1^[(μ ± zsμ)c] for coverage and exp[(β ± zsβ)c] for odds ratios, where μ and β are the maximum likelihood estimates of the log odds of coverage and community:team log odds ratio respectively, sμ and sβ are their standard errors, z is the 97.5% quantile of the standard normal distribution, and c is a factor that approximately corrects for bias when transforming GLMM estimates from log odds and log odds ratios to probability and odds ratios respectively^34,35^; c = √{1 + [16√3/(15π)]^2^}^-1^, where V is the sum of the GLMM random effect variance estimates.

### Outcome measure

The outcome measure is vaccination coverage recorded at each of the six survey visits. Vaccination coverage is defined as the count of dogs with a fully or partially completed vaccination certificate divided by the sum of the following counts: dogs with fully or partially completed vaccination certificates, and unvaccinated dogs (dogs without vaccination certificates). Because vaccination could not be verified, dogs claimed by their owner to be vaccinated but where a vaccination certificate could not be produced were excluded from both numerator and denominator. In order to assess the sensitivity of the trial results to excluding these dogs, the primary analysis (described below) was repeated using a more stringent definition of vaccination coverage where dogs stated by the owner to be vaccinated but without a certificate were considered unvaccinated and are therefore added to the denominator.

### Primary analysis

Mean coverage. The primary analysis was a test of the null hypothesis of equal coverage (community:team odds ratio = 1) between team-based and community-based delivery against the two-sided alternative hypothesis of unequal coverage. The full model, including the main effect of trial arm and the arm × year, arm × visit, and arm × year × visit interactions, was compared with a null model not including trial arm or its interactions with visit or year. In effect, the null model allows coverage to vary over the six time points (the six household surveys), while the full model includes an additional six parameters that allow coverage to vary independently between trial arms at each of the six time points. Under the null hypothesis, therefore, there is no intervention effect at any of the six time points, while the alternative hypothesis is that there is an intervention effect at one or more time points.

### Secondary analyses

Mean coverage at the eleven-month time point. Vaccination coverage at the second survey visit within each annual vaccination cycle was compared between arms to test the hypothesis that eleven-month vaccination coverage in community-based delivery was higher than the coverage at the same time-point in team-based delivery. This hypothesis was tested using the primary analysis model.

The probability of coverage within a village dipping below the critical vaccination threshold. The probability of coverage dipping below 40% was estimated in each arm. An overall estimate and an estimate at the midpoint of each month throughout the annual vaccination cycle (1st November to 31st October) were made. The probability distribution of coverage across wards by month (1-12) was estimated in each arm as follows: the primary analysis model was refitted, replacing the categorical variable survey visit by survey visit date; variation in model parameter estimates was estimated via 10,000 parametric bootstrap samples; for each bootstrap sample, coverage at each month was predicted assuming a logit-linear trend across each year within each arm; and random variation in coverage among wards, districts and survey visits was added by simulating 112 new wards per arm from the estimated random effect variances. The proportion of simulated coverages below the 40% threshold and 95% CI were estimated in each arm (overall and by month) as the mean, 2.5% and 97.5% centiles of the bootstrap samples.

Temporal variation in coverage. A key aim of Community-based MDV is greater consistency of vaccination coverage over time, and in particular to prevent coverage falling below the 40% threshold towards the end of the annual vaccination cycle. We compared three aspects of temporal variation in vaccination coverage between the two trial arms.

1. Consistency of coverage over the trial duration. We tested the hypothesis that mean vaccination coverage was more consistent over the three-years in the community-based arm than in the team-based arm. The mean absolute difference between the six consecutive time-points in the log odds of predicted coverage was estimated and exponentiated to an odds ratio, for each arm, and the community:team ratio of these inter-survey variation odds ratios calculated. A 95% CI for this ratio was estimated using 10,000 parametric bootstrap samples. Greater consistency over time through Community-based delivery would be indicated by a 95% CI below 1, while a 95% CI greater than 1 would indicate that Team-based delivery was more consistent.
2. Change in coverage from Survey Visit 1 to Visit 2. We hypothesised that vaccination coverage might fall from Survey Visit 1 to 2 in the Team-based arm while remaining relatively stable in the Community-based arm. We therefore estimated the Visit 1 to 2 coverage change odds ratio in each arm with Wald 95% CIs, and tested the null hypothesis that these odds ratios did not differ between arms, which is a test of the arm × visit interaction. To justify performing a single analysis over the whole trial duration rather than separately for each year, we first tested whether the arm × visit interaction was consistent over the three years (arm × visit × year interaction).
3. Minimum coverage at the start of the year required to ensure that coverage after one year does not drop below the critical vaccination threshold. The arm-specific Visit 1-2 odds ratios and 95% CIs calculated above, which cover a seven-month period, were used to extrapolate backwards from 40% end-of-year coverage to the required start-of-year coverage. Given that no vaccination took place after Visit 1 in the Team-based arm, the observed rate of change of coverage in this arm from Visit 1 to 2 is an estimate of the natural decay rate of vaccination coverage which occurs due to the expected death of vaccinated dogs and birth of unvaccinated puppies.

#### Inter-ward variation in coverage

We tested the hypothesis that community-based vaccination leads to greater spatial consistency in coverage by comparing inter-ward coverage variation between the two arms. The primary analysis model was modified to replace the single inter-ward random effect variance with two inter-ward variances, one per arm. This model was fitted using MCMC because inference on variance components is more reliable than using maximum likelihood. A 95% credible interval (CrI) from the posterior distribution of the community:team ratio of inter-ward variances was estimated. The null hypothesis of equal variances would be rejected if the 95% CrI for the ratio of variances did not include 1, with community-based delivery being considered more consistent than team-based if the 95% CrI were below 1.

### Sample size calculation

We estimated that randomizing 56 wards to each arm of the trial would give 87% power at the 5% significance level to detect a difference in mean coverage between team- and community-based MDV delivery, assuming mean coverage (the mean of coverage at 2 and 11 months) of 50% with team-based and 58% with community-based delivery (equivalent to an odds ratio of 1.34). Mean coverage at 2 months in both arms was assumed to be 60% following static-point vaccination clinics. Mean coverage is assumed to decline in the team-based arm to 41% at 11 months due to deaths and births (assuming an exponentially distributed lifespan with a mean of 26 months^31^, and is assumed to decline less sharply to 55% in the community-based arm due to ongoing vaccination. Power was estimated by analysing 10,000 simulated data sets, assuming sampling of one village per ward, three sub-villages per village, 10 households per sub-village and an average of 2.5 dogs per dog-owning household^31^ at 2 and 11 months for three years. Under this design, on average 8,400 dogs would be sampled at each of six timepoints, therefore 50,400 dogs would be expected to be surveyed in total. Logit-normal variances between households (11.5), sub-villages (4.4), villages (0.55) and wards (0.023) and the distribution of dogs over households were based on survey data from previous studies^5,36^, and have an effect on required sample size equivalent to a design effect of 23.

## Results

### Programmatic Results

The characteristics of the districts and randomised wards, overall and by trial arm, are given in Table In total 186,931 dogs were vaccinated during the three years and 25,677 dogs had their vaccination status confirmed through surveys (Figure 2). We implemented 99% of planned surveys in both trial arms, with 95% of surveys reaching at least 20 dog-owning households, recording a mean of 27 dogs per survey. On average 1,669 dogs were vaccinated in each of the 112 wards, with a mean of 1,050 and 2,289 dogs vaccinated in the team-based versus community-based arms, respectively. Vaccination coverage by ward and trial arm over the six time points is shown in Figures S1 and S2.

**Figure 2.**
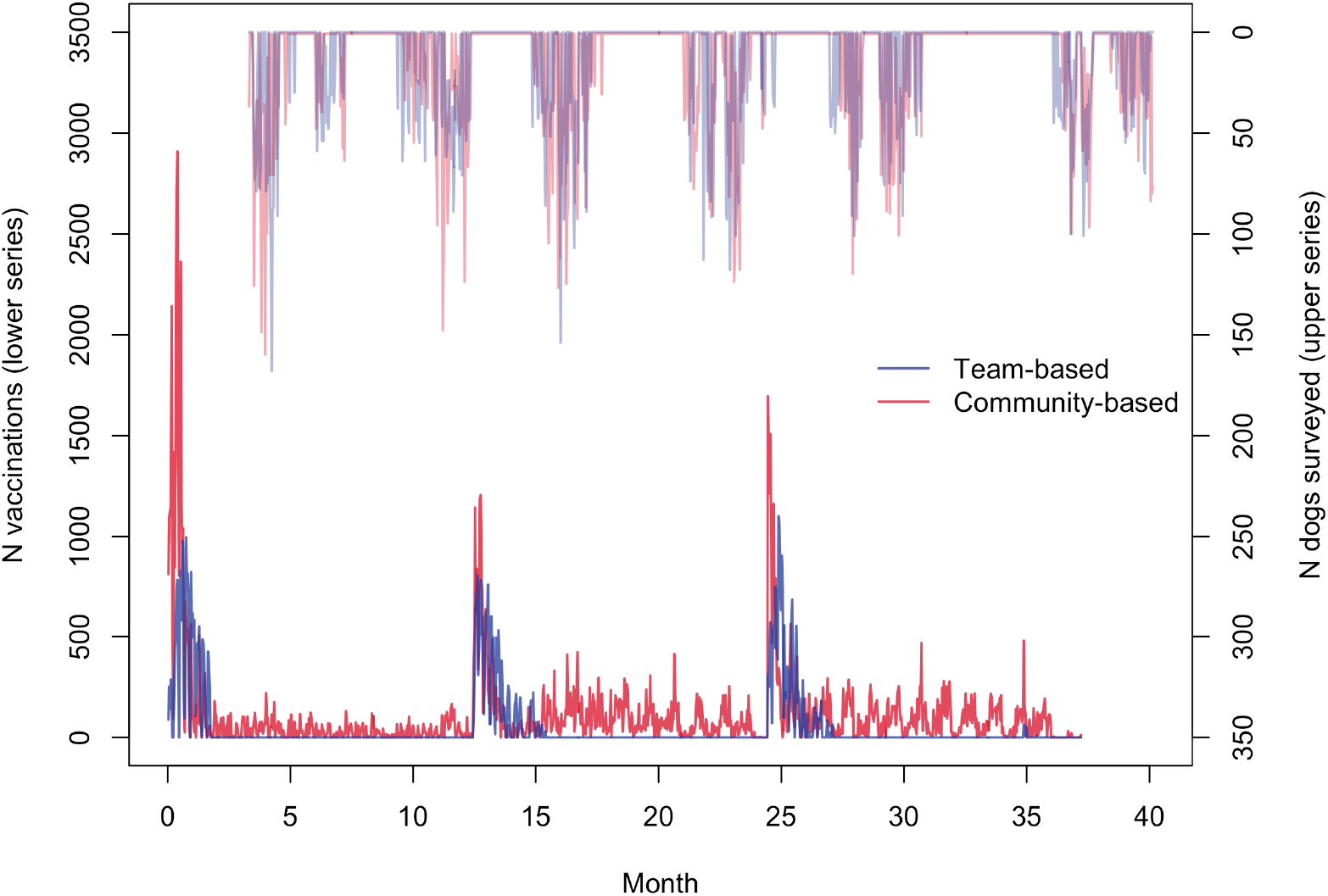
Number of rabies vaccinations performed (lower series; total = 186,931) and number of dogs surveyed (upper series; total = 25,677) per day by trial arm.

### Primary analysis

#### Mean coverage

The mean inter-campaign vaccination coverage for each trial arm is given in Table 2. Mean coverage differed between arms across the six time-points (P < 0.001), with mean coverages for each timepoint ranging from 22% to 46% in the team-based arm, and 49% to 62% in the community-based arm (Figure 3). Mean coverages from the first survey each year (month 1) ranged from 32% to 46% in the team-based arm, and 50% to 62% in the community-based arm, whilst in second surveys ranged from 22% to 31% in the team-based arm, and 49% to 57% in the community-based arm. The odds ratio estimates for the coverage difference reflected significantly higher coverage in the community-based arm at all six time points, ranging from 1.48 to 3.49 (Figure 4). Table S2 reports more detailed estimates from the primary analysis model. When the primary analysis was repeated using the more stringent definition of coverage, the between-arm comparison gave similar results, with coverage across the six surveys higher in the community-based arm (P < 0.001), although coverage estimates were substantially lower (Tables S3, S4; Figures S3, S4).

**Table 1.** Characteristics of randomised wards, overall and by trial arm.

|  |  | Total<br>(n=112) | Trial Arm |  |
| --- | --- | --- | --- | --- |
|  |  |  | Team-based<br>(N=56) | Community-based<br>(N=56) |
| District | Bunda | 15 (13%) | 7 (12%) | 8 (14%) |
|  | Butiama | 14 (12%) | 7 (12%) | 7 (12%) |
|  | Musoma | 28 (25%) | 15 (27%) | 13 (23%) |
|  | Rorya | 17 (15%) | 8 (14%) | 9 (16%) |
|  | Serengeti | 13 (12%) | 7 (12%) | 6 (11%) |
|  | Tarime | 25 (22%) | 12 (21%) | 13 (23%) |
| Land use | Rural | 83 (74%) | 37 (66%) | 46 (82%) |
|  | Semi-Urban | 7 (6%) | 5 (9%) | 2 (4%) |
|  | Urban | 22 (20%) | 14 (25%) | 8 (14%) |
| N dogs surveyed | Mean (SD) | 164 (55) | 165 (49) | 163 (61) |
|  | Median (IQR) | 162 (122, 192) | 171 (122, 190) | 159 (122, 192) |
|  | [Range] | [49, 444] | [76, 301] | [49, 444] |
| Survey completeness<br>(%) | Mean (SD) | 98.7 (5.1) | 98.2 (6.1) | 99.1 (3.8) |
|  | Median (IQR) | 100.0 (100.0, 100.0) | 100.0 (100.0, 100.0) | 100.0 (100.0, 100.0) |
|  | [Range] | [66.7, 100.0] | [66.7, 100.0] | [83.3, 100.0] |
| N dogs (estimated) | Mean (SD) | 2466 (1082) | 2398 (1142) | 2534 (1024) |
|  | Median (IQR) | 2420 (1804, 2954) | 2280 (1723, 2954) | 2493 (2057, 2933) |
|  | [Range] | [143, 6634] | [339, 6634] | [143, 5522] |
| N vaccinations over<br>3 years | Mean (SD) | 1669 (961) | 1050 (484) | 2289 (920) |
|  | Median (IQR) | 1584 (957, 2198) | 1002 (676, 1305) | 2217 (1756, 2684) |
|  | [Range] | [139, 5160] | [167, 2096] | [139, 5160] |

**Table 2.** Coverage and intervention odds ratio estimates (95% CI) at each survey timepoint estimated from the primary analysis GLMM (N dogs surveyed = 18358).

| Year-visit | Coverage (95% CI) |  | Odds ratio (95% CI) | P-value |
| --- | --- | --- | --- | --- |
|  | Team-based | Community-based |  |  |
| Y1-V1 | 0.32 (0.25, 0.41) | 0.53 (0.44, 0.62) | 2.36 (1.74, 3.20) | <0.001 |
| Y1-V2 | 0.22 (0.16, 0.29) | 0.49 (0.40, 0.58) | 3.49 (2.53, 4.81) | <0.001 |
| Y2-V1 | 0.40 (0.32, 0.49) | 0.50 (0.41, 0.58) | 1.48 (1.10, 2.01) | 0.011 |
| Y2-V2 | 0.31 (0.24, 0.40) | 0.57 (0.48, 0.65) | 2.90 (2.13, 3.93) | <0.001 |
| Y3-V1 | 0.46 (0.37, 0.54) | 0.62 (0.53, 0.70) | 1.96 (1.45, 2.66) | <0.001 |
| Y3-V2 | 0.31 (0.24, 0.39) | 0.54 (0.45, 0.63) | 2.62 (1.93, 3.56) | <0.001 |

**Figure 3.**
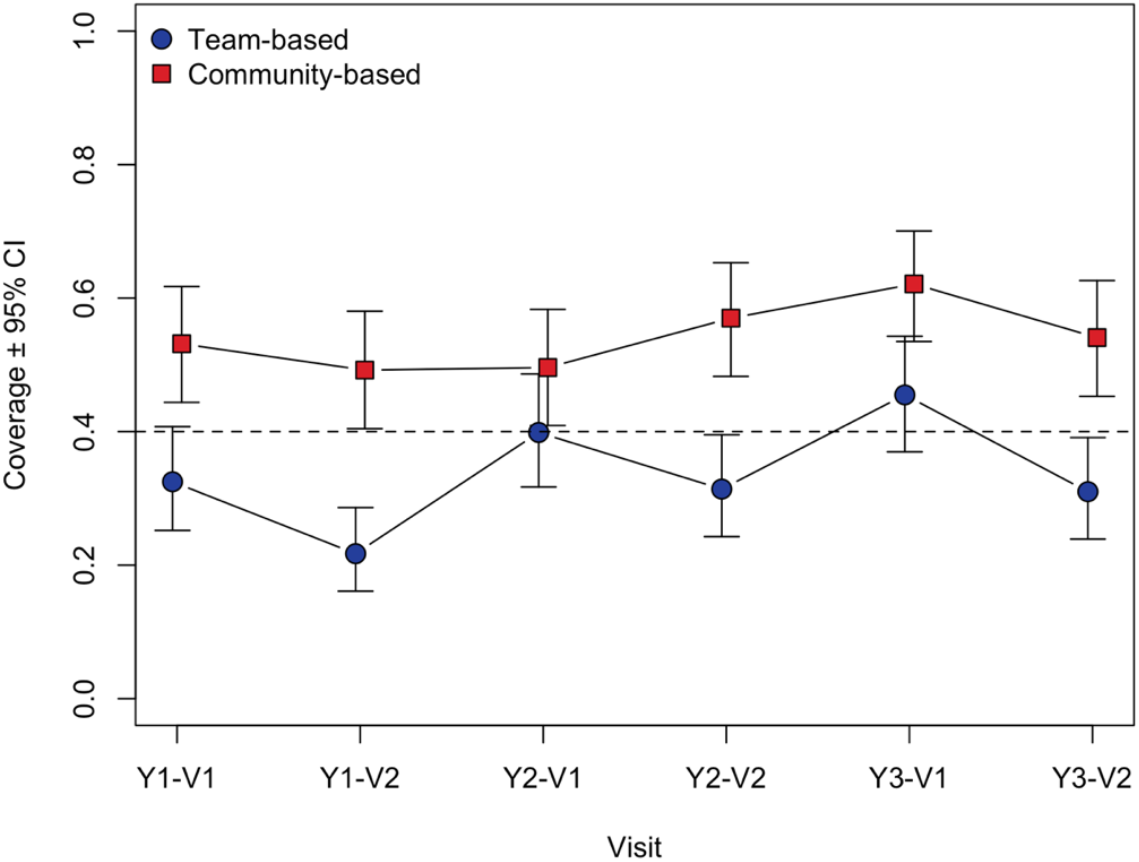
Estimated coverage ± 95% confidence limits at each survey time point, by trial arm.

**Figure 4.**
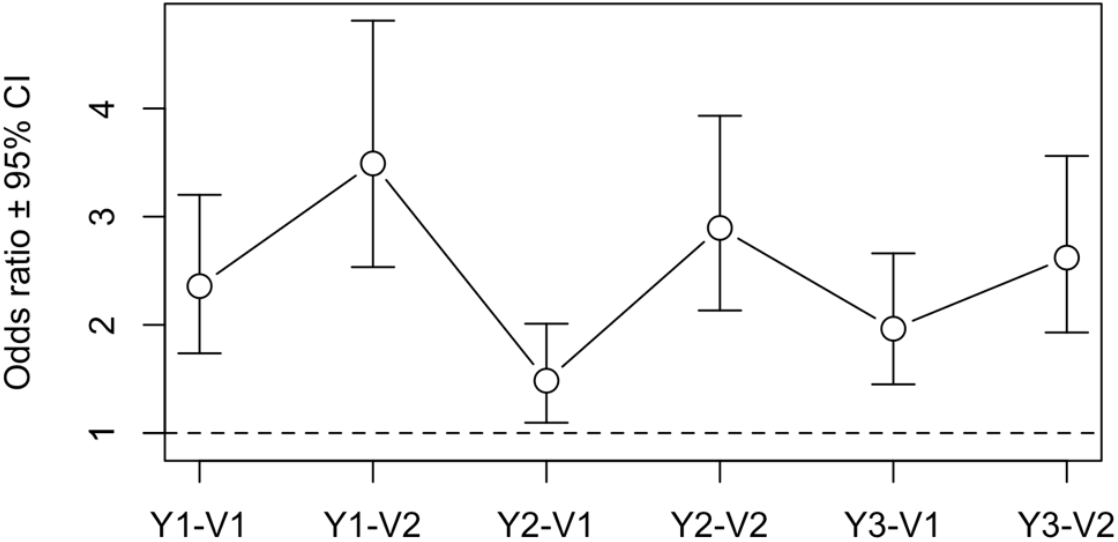
Estimated intervention effect (community:team) odds ratios ± 95% confidence limits at each survey time point.

### Secondary Analyses

#### Mean coverage during second survey

Mean coverage measured during the second surveys was consistently higher in community-based arm compared to the team-based arm (P < 0.001; Table 2). *The probability of coverage within a village dipping below the critical vaccination threshold*. Figure 5 shows the probability distribution of coverage in each arm across months 1 to 12 of the annual vaccination cycle. At the year start, coverage was generally higher in the community-based arm (month 1 mean: 57%) than the team-based arm (month 1 mean: 46%). As the year progresses, the coverage distribution remains stable in the community-based arm (month 12 mean: 53%) while dropping steadily in the team-based arm (month 12 mean: 29%). As a consequence, in the community-based arm the probability of a village falling below the 40% threshold is low and stable over time, from 20% in month 1 to 21% in month 12 (Table 3). By contrast, in the team-based arm the probability of a village falling below the 40% threshold is substantially higher than in the community-based arm in month 1 at 40%, and increases markedly across the year such that by the final month, 80% of villages are expected to fall below the critical coverage threshold. Averaged across the trial year, the probability of coverage within a village dipping below the critical vaccination threshold was 60% (95% CI: 38, 81) in the team-based arm, considerably higher than in the community-based arm at 18% (95% CI: 4, 40).

**Table 3.** Estimated probability (95% CI) of village-level coverage below 40% by month and averaged over the trial year, for each trial arm. The distribution of coverage among villages was estimated from 10,000 parametric bootstrap samples.

| Trial month | P(Coverage<0.4) (95% CI) |  |
| --- | --- | --- |
|  | Team-based | Community-based |
| 1 | 0.40 (0.19, 0.62) | 0.20 (0.05, 0.41) |
| 2 | 0.43 (0.21, 0.66) | 0.19 (0.05, 0.40) |
| 3 | 0.46 (0.23, 0.70) | 0.19 (0.04, 0.40) |
| 4 | 0.50 (0.26, 0.74) | 0.18 (0.04, 0.40) |
| 5 | 0.54 (0.30, 0.78) | 0.17 (0.03, 0.40) |
| 6 | 0.59 (0.33, 0.82) | 0.17 (0.02, 0.40) |
| 7 | 0.63 (0.38, 0.86) | 0.17 (0.02, 0.40) |
| 8 | 0.67 (0.42, 0.89) | 0.17 (0.02, 0.40) |
| 9 | 0.71 (0.46, 0.91) | 0.18 (0.02, 0.41) |
| 10 | 0.75 (0.51, 0.93) | 0.18 (0.03, 0.42) |
| 11 | 0.77 (0.54, 0.94) | 0.20 (0.04, 0.43) |
| 12 | 0.80 (0.57, 0.95) | 0.21 (0.05, 0.45) |
| All months | 0.60 (0.38, 0.81) | 0.18 (0.04, 0.40) |

**Figure 5.**
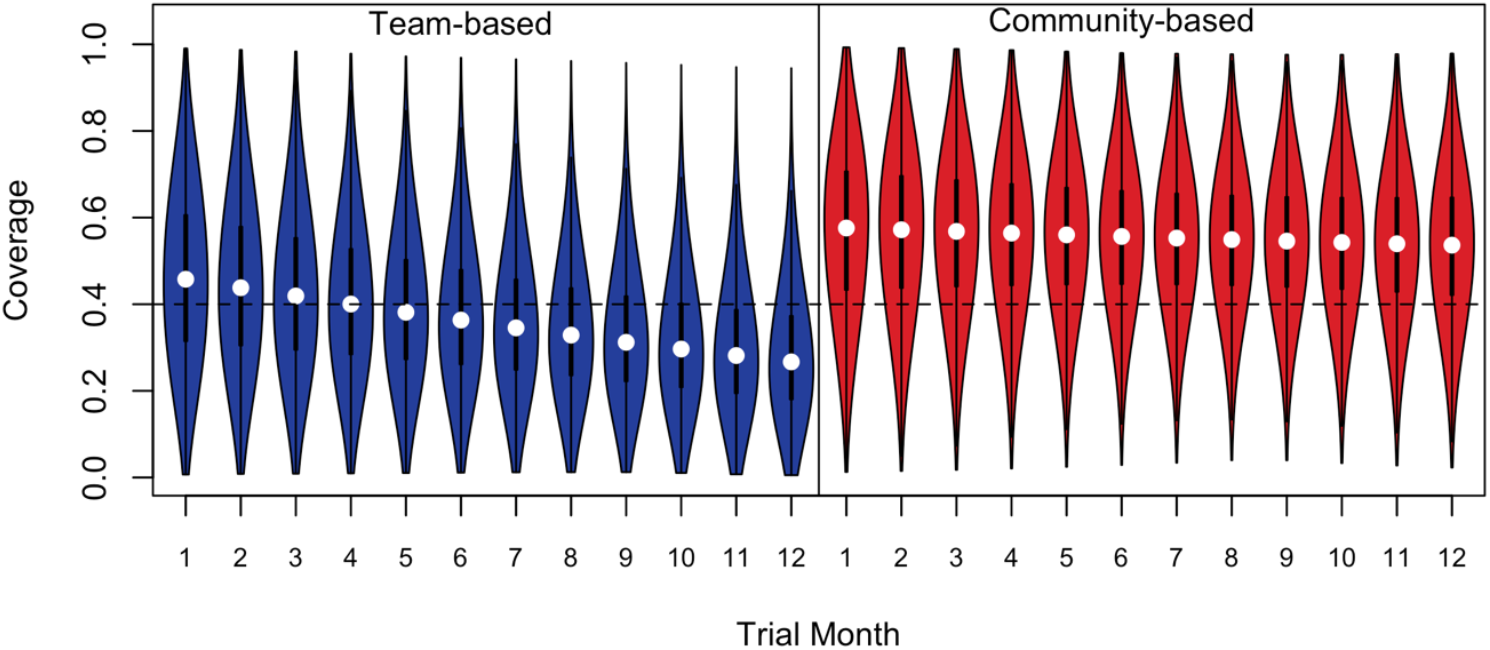
Estimated probability distribution of coverage by trial month for each arm. The coverage target of 40% is shown by a dashed line. White circles indicate medians and quartiles and range are shown by box-and-whisker plots.

Temporal variation in coverage:

1. *Consistency of coverage over the trial duration*. Variation in coverage over the six surveys, was lower in the Community-based arm (inter-survey odds ratio, i.e. community:team 0.67 [95% CI: 0.56, 0.83]).
2. *Inter-survey change in coverage*. Coverage fell from survey visit 1 to 2 in the team-based arm (odds ratio: 0.60 [95% CI: 0.50, 0.72]; P < 0.001), but not in the community-based arm (odds ratio: 0.94 [95% CI: 0.79, 1.13]; P = 0.533). The Visit 1:Visit 2 odds ratio was 1.58 (95% CI: 1.30, 1.92) times higher in the community-based arm than the team-based arm (arm × visit interaction P < 0.001). This interaction odds ratio can be interpreted as the community:team odds ratio being 1.58 times higher at Visit 2 than at Visit 1. The arm × visit interaction effect did not differ between the three years (arm × visit × year interaction P = 0.256), justifying the estimation and testing of a single interaction effect rather than three year-specific effects.
3. *Minimum coverage at the start of the year required to ensure that coverage after one year does not drop below the critical vaccination threshold*. The target coverage required at the year start to maintain coverage above 40% until the year end was 61% (95% CI: 53, 68) (Table 4) for team-based delivery versus 42% (95% CI: 35, 50) for community-based delivery, reflecting the relative stability of coverage over time in this arm.

**Table 4.** Estimate (95% CI) of Visit 2:Visit 1 odds ratios and of minimum start-of-year coverage required for end-of-year coverage to exceed 40%, by trial arm. The Visit 2:Visit 1 odds ratios differed significantly between trial arms (interaction P < 0.001).

| Trial Arm | Visit effect OR (95% CI) | P-value | Year beginning coverage for year end coverage ≥ threshold |
| --- | --- | --- | --- |
| Team-based | 0.60 (0.50, 0.72) | <0.001 | 0.61 (0.53, 0.68) |
| Community-based | 0.94 (0.79, 1.13) | 0.533 | 0.42 (0.35, 0.50) |

#### Inter-ward coverage variation

The community:team ratio of inter-ward variances was 2.21 (95% CrI: 0.09, 17.74); with no evidence for a difference in the amount of spatial variation in coverage. However, the wide 95% CrI indicates insufficient power to detect even substantial differences.

## Discussion

Our key findings were that: i. Rabies vaccination coverage in the community-based arm was consistently higher than that in the team-based arm; ii. The probability of vaccination coverage dropping below the critical vaccination threshold, below which herd immunity is lost, at any point in an annual cycle was much higher in the team-based arm, where coverage was below the critical threshold in more than 50% of villages during the latter part of the year (months five to 12); iii. initial coverage achieved needs to be much higher in team-based (61%) compared to community-based (42%) delivery to ensure coverage does not dip below the critical threshold by the year end.

### Critical vaccination threshold

The importance of keeping vaccination coverage above the critical vaccination threshold is well understood. Impacted by the relatively low R0 of rabies, which in most settings is approximately 1.1 - 1.2, the critical vaccination coverage threshold has been estimated to be between 20% and 40%^37^. To keep coverage above the upper bound (40%), the target coverage needed in an annual delivery approach has been estimated around 70%. Achieving this coverage at scale across the rural landscapes in rabies endemic settings is logistically challenging. We measured the vaccination decay rate to estimate what initial vaccination coverage would need to be to ensure it did not drop below the critical threshold. The resulting figure (61%) is lower than estimated previously^38^, but 19% higher than the target needed to be reached at the start of each annual cycle through continuous delivery via the community-based approach. Since vaccination occurs throughout the year, herd immunity is maintained in the community-based approach despite initial vaccination activity not reaching the 70% coverage target. Additionally, dog owners sometimes do not attend mass dog vaccination clinics because their dogs are not at home or cannot be caught or brought on the vaccination day, or because of scheduling conflicts with activities such as market days, planting seasons etc^5^. Having additional dates during the year when vaccination is carried out provides dog owners with more opportunity to vaccinate their dogs.

### Benefits of increasing coverage

Achieving a coverage greater than the critical vaccination coverage is important to control transmission and eliminate rabies locally. However, this does not mean that further increasing the coverage above the critical threshold is without merit and, when implementing vaccination campaigns, achieving the highest possible coverage is important for several reasons. First, as the number of immune individuals increases so does the speed with which transmission events die out and rabies control can be achieved. Second, as a result of more immune dogs and reduced transmission, the probability of a person being bitten declines, resulting in fewer people suffering the trauma of exposure to a suspect rabid dog, fewer people needing to urgently seek expensive PEP, and fewer people dying of rabies. For every additional percentage point of coverage achieved, therefore, numerous benefits of vaccination accrue. Given the demonstration that community-based delivery achieved a coverage considerably higher than team-based delivery, it seems likely that substantial and meaningful benefits can be realized, both at individual and national levels, further study investigating the broader benefits as well as economics and sustainability of community-based rabies control is warranted.

### Community driven delivery

International aid organizations and national governments increasingly favor bottom-up approaches^39^ that involve communities in the design and implementation of sustainable interventions^40^. Community-driven interventions have been instrumental in the fight against neglected tropical diseases. For example, community distributors of medicines for control of onchocerciasis have been involved in distributing medicines for up to 18 years, often without remuneration, demonstrating communities ownership of their health needs beyond financial incentives. In addition to normative benefits, community participation often leads to better outcomes, reduced costs, improved maintenance, efficiency, and enhanced social capital^41^. But questions remain about the level to which communities are willing to share the burden of these interventions and, consequently, the level of external support required to maintain the effectiveness of disease control activities. The problem of program sustainability has broader relevance regarding how donor funded public health initiatives can be maintained and how local community involvement impacts sustainability and equity. This has a particular relevance for dog-mediated human rabies because, although dog bites tend to occur very locally, dogs rarely (< 11%) bite a member of their own household (unpublished data), suggesting the need for community-level action. In this regard, rabies despite being maintained by animals, is like human infectious diseases where vaccination benefits are derived at the community-level through herd immunity effects.

### Thermotolerance

An important component of the community-based MDV was the requirement for vaccines to be stored within, and be available to, local communities throughout the year. For this to be possible in remote places where power resources and refrigeration units are scarce, the vaccines being used needed to be thermotolerant and retain potency following extended storage outside of the cold-chain. The Nobivac™ Rabies vaccine was determined to be thermotolerant^16,42^, with storage for up to 25°C for six months or 30°C for three months within these communities. To stabilize storage conditions as much as possible, vaccines were stored within each community inside a locally designed and tested passive cooling device, or Zeepot ^16^. The internal temperature of each Zeepot was recorded during the trial to ensure that the storage temperature did not exceed 30°C. In addition, any doses of vaccine that remained unused after six months, or three months if the temperature exceeded 25°C, were returned to the District Veterinary Office for disposal. In this way, the potency of the vaccines used in the trial was ensured.

The importance of cold-chain independent vaccine formulations in disease control was established in the 1990’s with the development of a thermostable rinderpest vaccine which was critical to the disease’s global eradication^43^. Establishing thermostability of other rabies vaccine formulations will be important if this key determinant of vaccination delivery to remote communities is to be further exploited to benefit rabies control. A consequence of having vaccines stored within communities was that they were available for use throughout the year, providing multiple opportunities to vaccinate dogs. For this system to work, dogs that had not been vaccinated earlier in each annual cycle needed to be identified, requiring an effective village-based OHC, selected because of their local knowledge and connectedness with the community. Consequently, the OHC had an important role in the success of the community-based approach, underscoring the importance of meaningful community engagement and involvement in community-driven initiatives.

We will in a subsequent paper compare the cost-effectiveness of these two delivery approaches in terms of cost per dog vaccinated and according to increased coverage attained, and therefore the economic implications for scaling that will depend on delivery costs. Similarly, our study took place in northern Tanzania across a region that is predominantly rural, with small areas of urbanisation. It is likely that the comparative assessment of the two approaches might differ in urban settings where dog ownership patterns might differ and where human demographic factors, such as population density, might impact the relationship that community vaccinators have and thereby their ability to reach dogs.

In conclusion, we demonstrate that community-based mass dog vaccination can achieve higher and more consistent coverage than the standard team-based approach. We do not suggest that the community-based approach will be suitable to all rabies-endemic regions. To achieve widespread rabies control a range of different, locally appropriate dog vaccination delivery methods will be required. For example, in countries with well-resourced animal health sectors, synchronised and nationally or regionally advertised single event mass dog vaccination approaches might be more effective at reaching more dogs at less cost. Nonetheless, the tested community-based approach has evident merit and can achieve consistently high coverage across a range of rural, peri-urban and urban settings broadly typical of many countries in sub-Saharan Africa. Gavi, the Vaccine alliance is now investing in human rabies post-exposure vaccines and requires eligible countries to develop comprehensive and sustainable dog vaccination programs. Our trial results suggest that community-based delivery strategies have an important role to play in these national rabies elimination strategies, and contribute to the global ‘zero by 30’ goal to end human rabies deaths by 2030^44^.

## References

1 Hampson K, Coudeville L, Lembo T, et al. Estimating the global burden of endemic canine rabies. PLoS Negl Trop Dis 2015; 9: e0003709.

2 Knobel DL, Cleaveland S, Coleman PG, et al. Re-evaluating the burden of rabies in Africa and Asia. Bull World Health Organ 2005; 83: 360–8.

3 Lankester F, Hampson K, Lembo T, Palmer G, Taylor L, Cleaveland S. Implementing Pasteur’s vision for rabies elimination. Science 2014; 345: 1562–4.

4 Minghui R, Stone M, Semedo MH, Nel L. New global strategic plan to eliminate dog-mediated rabies by 2030. Lancet Glob Health 2018; 6: e828–9.

5 Minyoo AB, Steinmetz M, Czupryna A, et al. Incentives Increase Participation in Mass Dog Rabies Vaccination Clinics and Methods of Coverage Estimation Are Assessed to Be Accurate. PLoS Negl Trop Dis 2015; 9: e0004221.

6 Taylor LH, Nel LH. Global epidemiology of canine rabies: past, present, and future prospects. Vet Med Res Rep 2015; : 361–71.

7 Cleaveland S, Hampson K. Rabies elimination research: juxtaposing optimism, pragmatism and realism. Proc R Soc B Biol Sci 2017; 284. DOI:10.1098/rspb.2017.1880.

8 Gibson AD, Handel IG, Shervell K, et al. The Vaccination of 35,000 Dogs in 20 Working Days Using Combined Static Point and Door-to-Door Methods in Blantyre, Malawi. PLoS Negl Trop Dis 2016; 10. DOI:10.1371/journal.pntd.0004824.

9 Coleman PG, Dye C. Immunization coverage required to prevent outbreaks of dog rabies. Vaccine 1996; 14: 185–6.

10 Ferguson EA, Hampson K, Cleaveland S, et al. Heterogeneity in the spread and control of infectious disease: consequences for the elimination of canine rabies. Sci Rep 2015; 5: 1–13.

11 Townsend SE, Sumantra IP, Pudjiatmoko, et al. Designing programs for eliminating canine rabies from islands: Bali, Indonesia as a case study. PLoS Negl Trop Dis 2013; 7: e2372.

12 Duamor CT, Hampson K, Lankester F, et al. Development, feasibility and potential effectiveness of community-based continuous mass dog vaccination delivery strategies: Lessons for optimization and replication. PLoS Negl Trop Dis 2022; 16: e0010318.

13 Mazeri S, Gibson AD, Meunier N, et al. Barriers of attendance to dog rabies static point vaccination clinics in Blantyre, Malawi. PLoS Negl Trop Dis 2018; 12: e0006159.

14 Hotez PJ, Fenwick A, Savioli L, Molyneux DH. Rescuing the bottom billion through control of neglected tropical diseases. The Lancet 2009; 373: 1570–5.

15 Karp CL, Lans D, Esparza J, et al. Evaluating the value proposition for improving vaccine thermostability to increase vaccine impact in low and middle-income countries. Vaccine 2015; 33: 3471–9.

16 Lugelo A, Hampson K, Ferguson EA, et al. Development of dog vaccination strategies to maintain herd immunity against rabies. Viruses 2022; 14: 830.

17 Zipursky S, Djingarey MH, Lodjo J-C, Olodo L, Tiendrebeogo S, Ronveaux O. Benefits of using vaccines out of the cold chain: delivering meningitis A vaccine in a controlled temperature chain during the mass immunization campaign in Benin. Vaccine 2014; 32: 1431–5.

18 Hipgrave DB, Maynard JE, Biggs B-A. Improving birth dose coverage of hepatitis B vaccine. Bull World Health Organ 2006; 84: 65–71.

19 Hipgrave DB, Tran TN, Huong VM, et al. Immunogenicity of a locally produced hepatitis B vaccine with the birth dose stored outside the cold chain in rural Vietnam. Am J Trop Med Hyg 2006; 74: 255–60.

20 Otto BF, Suarnawa IM, Stewart T, et al. At-birth immunisation against hepatitis B using a novel pre-filled immunisation device stored outside the cold chain. Vaccine 1999; 18: 498–502.

21 Sutanto A, Suarnawa I, Nelson C, Stewart T, Soewarso TI. Home delivery of heat-stable vaccines in Indonesia: outreach immunization with a prefilled, single-use injection device. Bull World Health Organ 1999; 77: 119.

22 Wang L, Li J, Chen H, et al. Hepatitis B vaccination of newborn infants in rural China: evaluation of a village-based, out-of-cold-chain delivery strategy. Bull World Health Organ 2007; 85: 688– 94.

23 Mgomezulu R, Alders R, Chikungwa P, Young M, Lipita W, Wanda G. Trials with a thermotolerant I-2 Newcastle disease vaccine in confined Australorp chickens and scavenging village chickens in Malawi. 2021; published online Jan 4.

24 Henderson D, Klepac P. Lessons from the eradication of smallpox: an interview with DA Henderson. Philos Trans R Soc B Biol Sci 2013; 368: 20130113.

25 Mariner JC, House JA, Mebus CA, et al. Rinderpest Eradication: Appropriate Technology and Social Innovations. Science 2013; 337: 1309–12.

26 Lugelo A, Hampson K, Czupryna A, et al. Investigating the Efficacy of a Canine Rabies Vaccine Following Storage Outside of the Cold-Chain in a Passive Cooling Device. Front Vet Sci 2021; 8: 1121.

27 Lugelo A, Hampson K, Bigambo M, Kazwala R, Lankester F. Controlling Human Rabies: The Development of an Effective, Inexpensive and Locally Made Passive Cooling Device for Storing Thermotolerant Animal Rabies Vaccines. Trop Med Infect Dis 2020; 5: 130.

28 Lankester FJ, Wouters P, Czupryna A, et al. Thermotolerance of an inactivated rabies vaccine for dogs. Vaccine 2016; 34: 5504–11.

29 Kaare M, Lembo T, Hampson K, et al. Rabies control in rural Africa: evaluating strategies for effective domestic dog vaccination. Vaccine 2009; 27: 152–60.

30 NBS. 2012 population and housing census. National Bureau of Statistics, 2012 https://www.nbs.go.tz/index.php/en/census-surveys/population-and-housing-census (accessed June 22, 2024).

31 Czupryna AM, Brown JS, Bigambo MA, et al. Ecology and demography of free-roaming domestic dogs in rural villages near Serengeti National Park in Tanzania. PLoS One 2016; 11: e0167092.

32 Hayes RJ, Moulton LH. Cluster randomised trials. Chapman and Hall/CRC, 2017 https://www.taylorfrancis.com/books/mono/10.4324/9781315370286/cluster-randomised-trials-richard-hayes-lawrence-moulton (accessed Oct 26, 2024).

33 R Development Core Team. R: A Language and Environment for Statistical Computing. 2024. https://www.R-project.org/.

34 Nakagawa S, Johnson PC, Schielzeth H. The coefficient of determination R 2 and intra-class correlation coefficient from generalized linear mixed-effects models revisited and expanded. J R Soc Interface 2017; 14: 20170213.

35 Zeger SL, Liang K-Y, Albert PS. Models for longitudinal data: a generalized estimating equation approach. Biometrics 1988; : 1049–60.

36 Sambo M, Johnson PCD, Hotopp K, et al. Comparing Methods of Assessing Dog Rabies Vaccination Coverage in Rural and Urban Communities in Tanzania. Front Vet Sci 2017; 4. DOI:10.3389/fvets.2017.00033.

37 Hampson KM, Chin SS, Mallen EA. Dual wavefront sensing channel monocular adaptive optics system for accommodation studies. Opt Express 2009; 17: 18229–40.

38 Hampson K, Dushoff J, Cleaveland S, et al. Transmission Dynamics and Prospects for the Elimination of Canine Rabies. PLoS Biol 2009; 7. DOI:10.1371/journal.pbio.1000053.

39 Labonne J, Chase RS. Do community-driven development projects enhance social capital? Evidence from the Philippines. J Dev Econ 2011; 96: 348–58.

40 Amazigo UV, Leak SGA, Zoure HGM, et al. Community-directed distributors—The “foot soldiers” in the fight to control and eliminate neglected tropical diseases. PLoS Negl Trop Dis 2021; 15: e0009088.

41 Chase R, Woolcock M. Social capital and the micro-institutional foundations of CDD approaches in East Asia: Evidence, theory, and policy implications. In: Arusha conference ‘New Frontiers of Social Policy’, December. 2005:12–5.

42 Lankester F, Lugelo A, Werling D, et al. The efficacy of alcelaphine herpesvirus-1 (AlHV-1) immunization with the adjuvants Emulsigen® and the monomeric TLR5 ligand FliC in zebu cattle against AlHV-1 malignant catarrhal fever induced by experimental virus challenge. Vet Microbiol 2016; 195: 144–53.

43 Mariner JC, House JA, Mebus CA, et al. Rinderpest eradication: appropriate technology and social innovations. Science 2012; 337: 1309–12.

44 WHO. Zero by 30: the global strategic plan to end human deaths from dog-mediated rabies by 2030. World Health Organization, 2018 https://www.who.int/rabies/resources/9789241513838/en/ (accessed April 20, 2021).

