## Supplementary material for "A randomized controlled trial of the effectiveness of a community-based rabies vaccination strategy": Figure S1

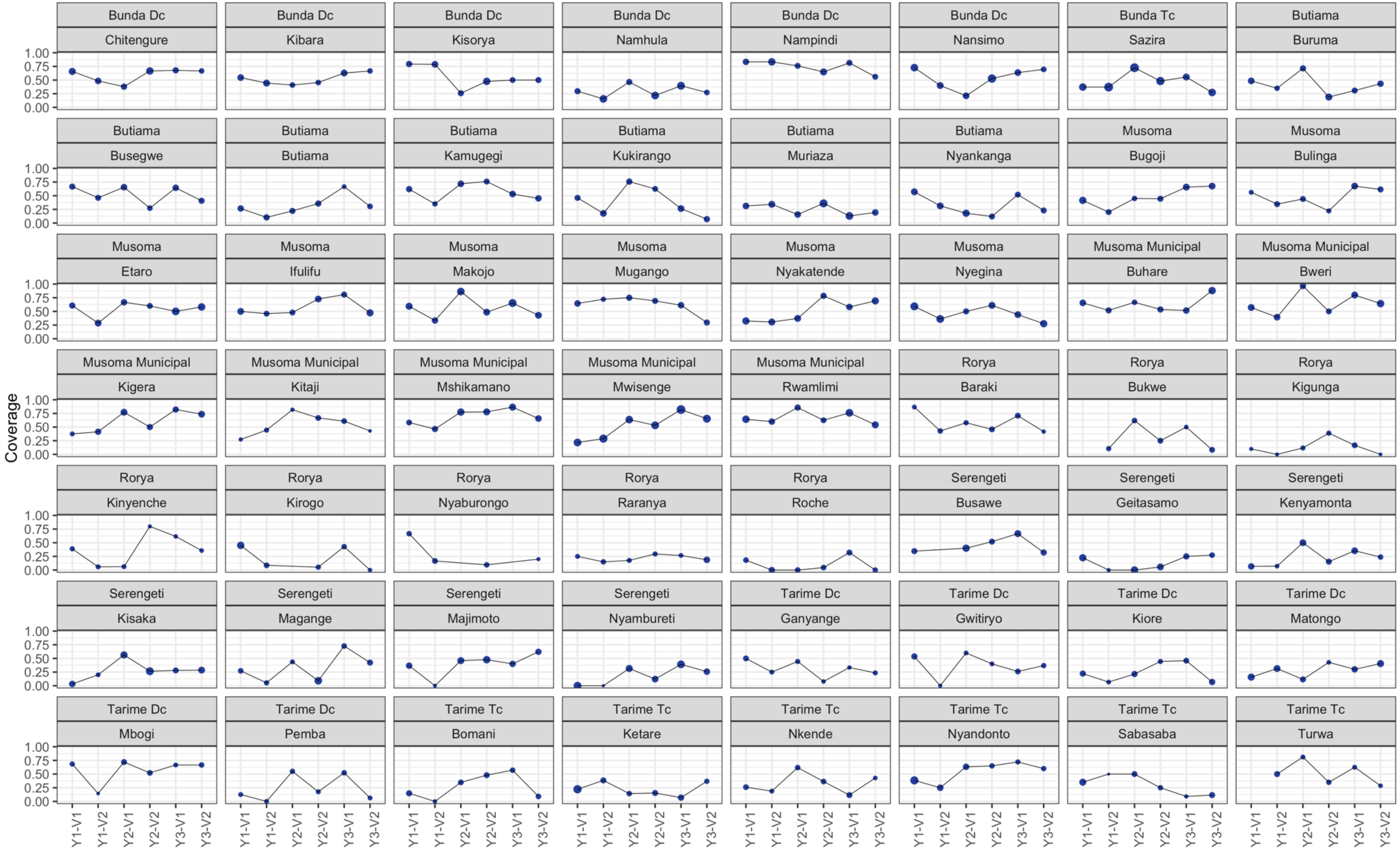


**Figure S1.** Coverage by ward and survey time point in the Team-based arm. Point area is proportional to the number of dogs surveyed.
