## Supplementary material for "A randomized controlled trial of the effectiveness of a community-based rabies vaccination strategy": Figure S2

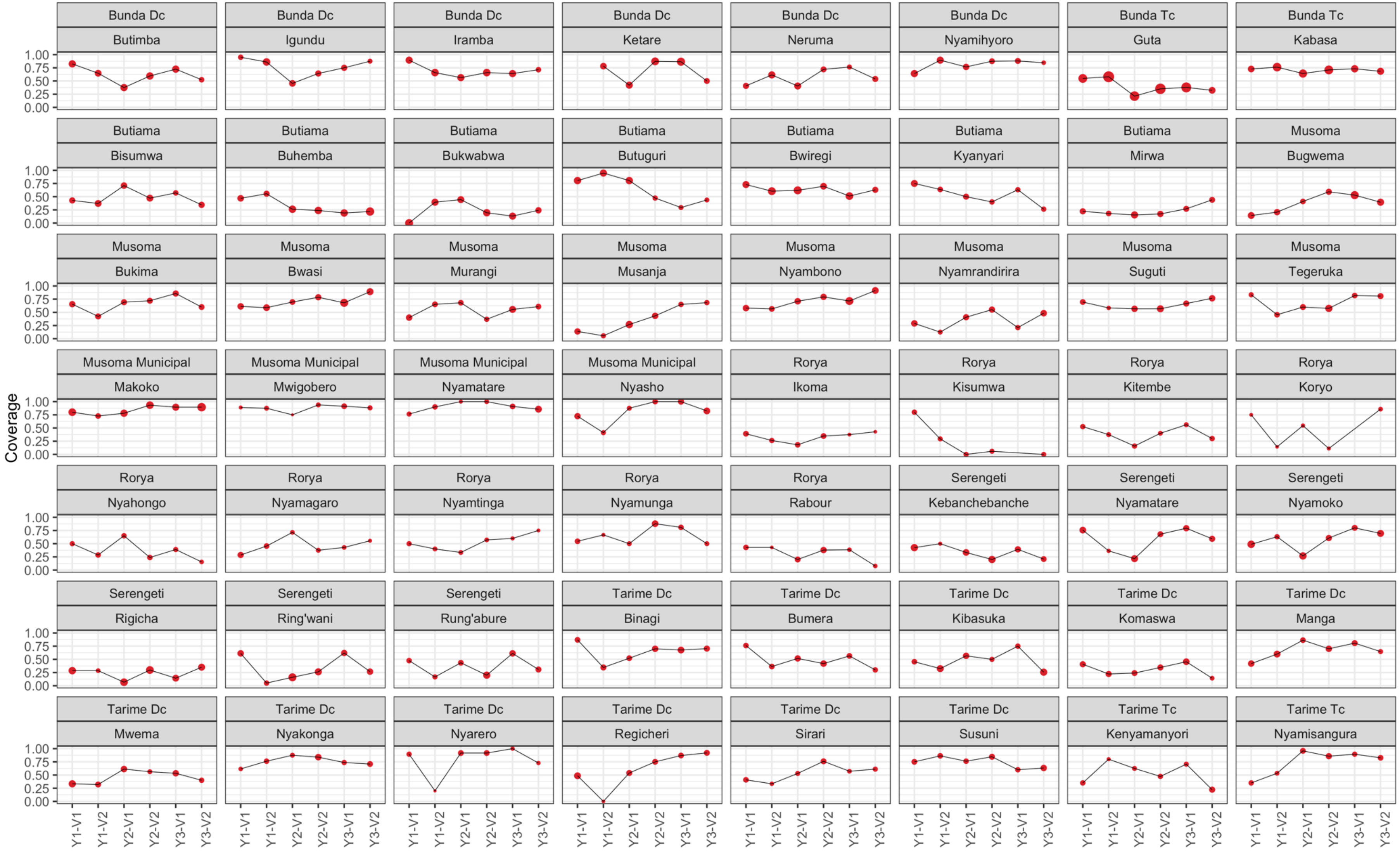


**Figure S2.** Coverage by ward and survey time point in the Community-based arm. Point area is proportional to the number of dogs surveyed.
