## Supplementary material for "A randomized controlled trial of the effectiveness of a community-based rabies vaccination strategy": Figure S3

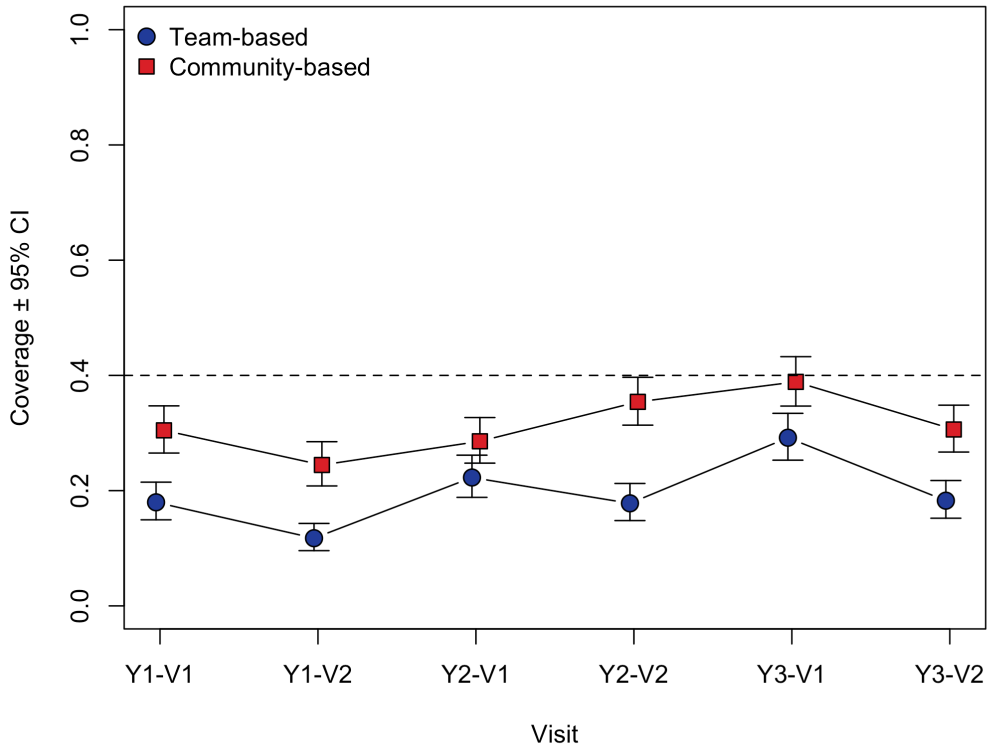


**Figure S3.** Estimated coverage ± 95% confidence limits at each survey time point, by trial arm. A more stringent definition of coverage was used, where dogs claimed to be vaccinated but where no vaccination certificate could be produced were assumed to be unvaccinated.
