## Supplementary material for "A randomized controlled trial of the effectiveness of a community-based rabies vaccination strategy": Figure S4

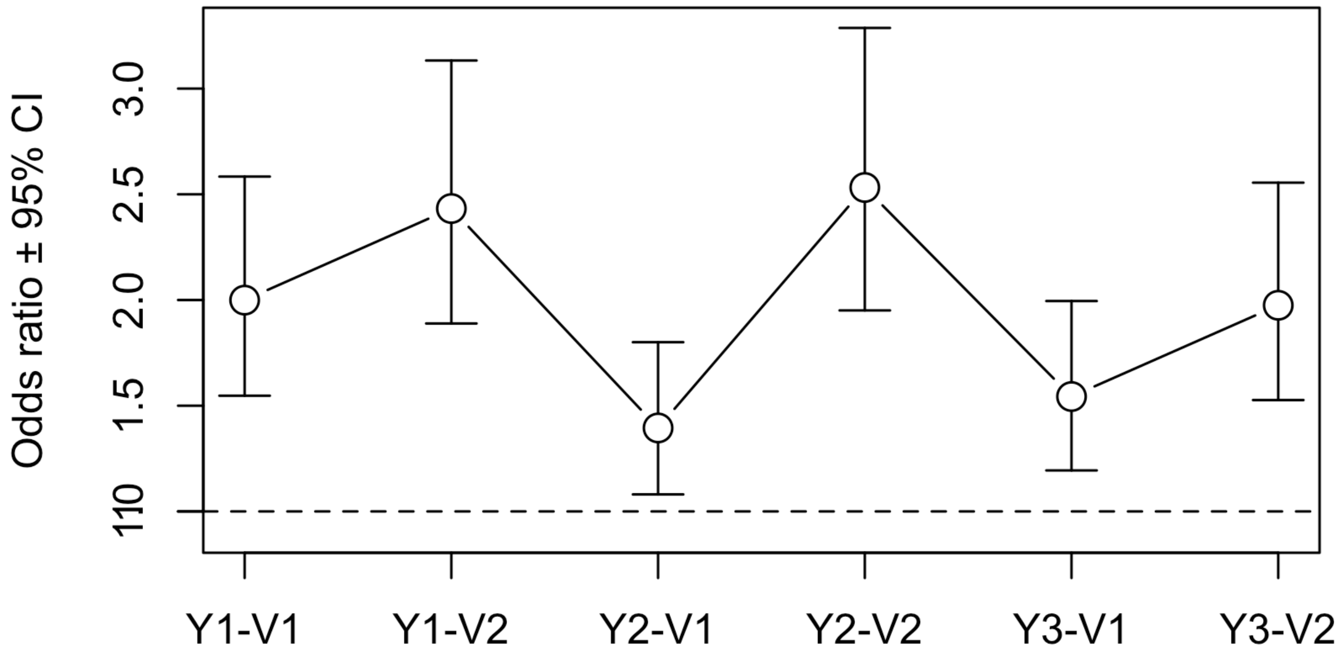


**Figure S4.** Estimated intervention effect (community:team) odds ratios ± 95% confidence limits at each survey time point. A more stringent definition of coverage was used, where dogs claimed to be vaccinated but where no vaccination certificate could be produced were assumed to be unvaccinated.
