## Supplementary material for "A randomized controlled trial of the effectiveness of a community-based rabies vaccination strategy": Table S1

**Table S1.** Summary of methods used for primary and secondary analyses. Generalized linear mixed-effects models (GLMMs) were fitted by maximum likelihood (ML) using the glmmTMB package and by Monte Carlo Markov-Chain (MCMC) using the brms package.

| Analysis | Question | GLMM fixed effects | GLMM random effects (n categories) | GLMM fitting method | Estimates |
|---|---|---|---|---|---|
| Primary 1 | Does mean coverage differ between arms at any of the six time-points? | arm + visit + year + arm × visit + arm × year + visit × year + arm × visit × year | District (9), ward (112), subvillage (659); visit-specific: district (54), ward (663); subvillage (2376); household (12999) | ML | Odds ratio estimate & Wald 95% CI at each survey visit in each year; LRT P-value for null hypothesis that each OR = 1. |
| Secondary 1 | Does coverage differ between arms at any of the three Survey Visit 2 time-points? | Same as Primary 1 | Same as Primary 1 | ML | Odds ratio estimate & Wald 95% CI at survey visit 2 in each year (already calculated for Primary analysis 1); LRT P-value for null hypothesis that each OR = 1. |
| Secondary 2 | What is the probability of coverage in a ward being below 40% in each arm, per trial month and averaged over the year. | arm + month + year + arm × month + arm × year | Same as Primary 1 | ML | Distribution of predicted coverage by arm and month estimated by simulating new wards, incorporating model uncertainty via 10,000 parametric bootstrap samples; P(coverage < 40%) by arm and month & 95% CI |
| Secondary 3.1 | Does the consistency of coverage over the duration of the trial differ between arms? | Same as Primary 1 | Same as Primary 1 | ML | Mean absolute difference between 6 consecutive time-points in log odds of predicted coverage exponentiated to an odds ratio, for each arm; the community:team ratio of odds ratios; |

| | | | | | |
|---|---|---|---|---|---|
| | | | | | 95% CI for each ratio, estimated using 10,000 parametric bootstrap samples |
| Secondary 3.2 | Does the rate of change in coverage from survey visit 1 to survey visit 2 differ between arms? | arm + visit + year + arm × visit + arm × year | Same as Primary 1 | ML | arm × visit interaction odds ratio, Wald 95% CI and Wald P-value; Survey Visit effect odds ratio, Wald 95% CI and Wald P-value, for each arm. |
| Secondary 3.3 | Given the observed rate of change of coverage within a year, what is the minimum coverage required at Survey Visit 1 to ensure that coverage at Survey Visit 2 does not drop below the critical vaccination threshold? | Same as Secondary 3.2 | Same as Primary 1 | ML | Coverage at survey visit 1 predicted to give 40% coverage at survey visit 2 & Wald 95% CI, calculated using the odds ratio estimates from Secondary Analysis 3.2. |
| Secondary 4 | Does the consistency of coverage over space (i.e. between wards) differ between arms? | Same as Primary 1 | Same as Primary 1, except that the inter-ward random effect variance varies between arms | MCMC | Median and 95% credible interval from the posterior distribution of the Community:Team ratio of inter-ward variances |
