## Supplementary material for "A randomized controlled trial of the effectiveness of a community-based rabies vaccination strategy": Table S2

**Table S2.** Estimates of fixed effects (log odds and log odds ratios) and random effects (variances) from the GLMMs fitted for the primary analysis. Numbers of observations, number of each random effect level, and marginal and conditional $R^2$ are also presented.

| | Primary analysis null hypothesis model | | | | Primary analysis alternative hypothesis model | | | |
|---|---|---|---|---|---|---|---|---|
| Predictors | Log-Odds | std. Error | CI | p | Log-Odds | std. Error | CI | p |
| (Intercept) | -0.60 | 0.32 | -1.23, 0.02 | 0.060 | -1.48 | 0.37 | -2.19, -0.76 | <0.001 |
| Visit: V 2 | -0.71 | 0.26 | -1.22, -0.21 | 0.006 | -1.11 | 0.32 | -1.73, -0.49 | <0.001 |
| Year: Y 2 | 0.16 | 0.26 | -0.35, 0.66 | 0.542 | 0.65 | 0.31 | 0.04, 1.25 | 0.036 |
| Year: Y 3 | 0.90 | 0.26 | 0.40, 1.41 | <0.001 | 1.11 | 0.31 | 0.51, 1.71 | <0.001 |
| VisitV2:YearY2 | 0.66 | 0.36 | -0.05, 1.38 | 0.068 | 0.36 | 0.44 | -0.50, 1.22 | 0.410 |
| VisitV2:YearY3 | -0.24 | 0.36 | -0.95, 0.48 | 0.514 | -0.14 | 0.44 | -1.00, 0.72 | 0.751 |
| Trial Arm: Community-based | | | | | 1.73 | 0.31 | 1.11, 2.35 | <0.001 |
| VisitV2:Trial_ArmCommunity-based | | | | | 0.79 | 0.35 | 0.10, 1.49 | 0.026 |
| YearY2:Trial_ArmCommunity-based | | | | | -0.93 | 0.34 | -1.61, -0.26 | 0.007 |
| YearY3:Trial_ArmCommunity-based | | | | | -0.37 | 0.34 | -1.04, 0.30 | 0.281 |
| VisitV2:YearY2:Trial_ArmCommunity-based | | | | | 0.56 | 0.49 | -0.40, 1.52 | 0.256 |
| VisitV2:YearY3:Trial_ArmCommunity-based | | | | | -0.21 | 0.49 | -1.17, 0.75 | 0.669 |
| Random Effects | | | | | | | | |
| $\sigma^2$ | 3.29 | | | | 3.29 | | | |
| $\tau_{00}$ | 3.68 $_{Household.file.id}$ | | | | 3.81 $_{Household.file.id}$ | | | |
| | 1.04 $_{sub\_village.rnd}$ | | | | 1.09 $_{sub\_village.rnd}$ | | | |
| | 2.30 $_{sub\_village}$ | | | | 2.36 $_{sub\_village}$ | | | |
| | 0.47 $_{ward.rnd}$ | | | | 0.35 $_{ward.rnd}$ | | | |
| | 1.25 $_{ward}$ | | | | 0.53 $_{ward}$ | | | |
| | 0.15 $_{district.rnd}$ | | | | 0.16 $_{district.rnd}$ | | | |
| | 0.46 $_{district}$ | | | | 0.58 $_{district}$ | | | |
| N | 12999 $_{Household.file.id}$ | | | | 12999 $_{Household.file.id}$ | | | |
| | 2376 $_{sub\_village.rnd}$ | | | | 2376 $_{sub\_village.rnd}$ | | | |
| | 659 $_{sub\_village}$ | | | | 659 $_{sub\_village}$ | | | |
| | 663 $_{ward.rnd}$ | | | | 663 $_{ward.rnd}$ | | | |
| | 112 $_{ward}$ | | | | 112 $_{ward}$ | | | |
| | 54 $_{district.rnd}$ | | | | 54 $_{district.rnd}$ | | | |
| | 9 $_{district}$ | | | | 9 $_{district}$ | | | |
| Observations | 18358 | | | | 18358 | | | |
| Marginal $R^2$ / Conditional $R^2$ | 0.017 / 0.744 | | | | 0.079 / 0.751 | | | |
