## Supplementary material for "A randomized controlled trial of the effectiveness of a community-based rabies vaccination strategy": Table S3

**Table S3.** Primary analysis (sensitivity analysis). Coverage and intervention odds ratio estimates (95% CI) at each survey estimated from the primary analysis GLMM re-fitted using a more stringent definition of coverage, where dogs claimed to be vaccinated but where no vaccination certificate could be produced were assumed to be unvaccinated (N dogs surveyed = 25677). The P-value is from a likelihood ratio test of the primary analysis null hypothesis of no intervention effect at any of the six surveys.

| Year-visit | Coverage (95% CI) | | Odds ratio (95% CI) | P-value |
|---|---|---|---|---|
| | Team | Community | | |
| Y1-V1 | 0.18 (0.15, 0.21) | 0.30 (0.27, 0.35) | 2.00 (1.55, 2.58) | <0.001 |
| Y1-V2 | 0.12 (0.10, 0.14) | 0.24 (0.21, 0.28) | 2.43 (1.89, 3.13) | <0.001 |
| Y2-V1 | 0.22 (0.19, 0.26) | 0.29 (0.25, 0.33) | 1.39 (1.08, 1.80) | 0.011 |
| Y2-V2 | 0.18 (0.15, 0.21) | 0.35 (0.31, 0.40) | 2.53 (1.95, 3.29) | <0.001 |
| Y3-V1 | 0.29 (0.25, 0.33) | 0.39 (0.35, 0.43) | 1.54 (1.19, 2.00) | 0.001 |
| Y3-V2 | 0.18 (0.15, 0.22) | 0.31 (0.27, 0.35) | 1.98 (1.53, 2.55) | <0.001 |
