## Supplementary material for "A randomized controlled trial of the effectiveness of a community-based rabies vaccination strategy": Table S4

**Table S4.** Estimates of fixed effects (log odds and log odds ratios) and random effects (variances) from the GLMMs fitted for the primary analysis. The primary analysis GLMMs were re-fitted using a more stringent definition of coverage, where dogs claimed to be vaccinated but where no vaccination certificate could be produced were assumed to be unvaccinated. Numbers of observations, number of each random effect level, and marginal and conditional $R^2$ are also presented.

| | Primary analysis null hypothesis model | | | | Primary analysis alternative hypothesis model | | | |
|---|---|---|---|---|---|---|---|---|
| Predictors | Log-Odds | std. Error | CI | p | Log-Odds | std. Error | CI | p |
| (Intercept) | -2.39 | 0.24 | -2.86, -1.91 | <0.001 | -3.96 | 0.29 | -4.53, -3.38 | <0.001 |
| Visit: V 2 | -0.85 | 0.30 | -1.43, -0.27 | 0.004 | -1.30 | 0.31 | -1.91, -0.69 | <0.001 |
| Year: Y 2 | 0.16 | 0.29 | -0.41, 0.73 | 0.580 | 0.70 | 0.32 | 0.08, 1.33 | 0.028 |
| Year: Y 3 | 0.95 | 0.29 | 0.37, 1.52 | 0.001 | 1.65 | 0.32 | 1.01, 2.28 | <0.001 |
| VisitV2:YearY2 | 0.89 | 0.41 | 0.07, 1.70 | 0.033 | 0.57 | 0.44 | -0.30, 1.44 | 0.199 |
| VisitV2:YearY3 | -0.15 | 0.41 | -0.96, 0.66 | 0.710 | -0.30 | 0.44 | -1.17, 0.58 | 0.506 |
| Trial Arm: Community-based | | | | | 1.81 | 0.34 | 1.14, 2.47 | <0.001 |
| VisitV2:Trial_ArmCommunity-based | | | | | 0.51 | 0.44 | -0.35, 1.37 | 0.245 |
| YearY2:Trial_ArmCommunity-based | | | | | -0.94 | 0.44 | -1.81, -0.07 | 0.034 |
| YearY3:Trial_ArmCommunity-based | | | | | -0.67 | 0.44 | -1.54, 0.19 | 0.127 |
| VisitV2:YearY2:Trial_ArmCommunity-based | | | | | 1.04 | 0.63 | -0.19, 2.27 | 0.096 |
| VisitV2:YearY3:Trial_ArmCommunity-based | | | | | 0.13 | 0.62 | -1.09, 1.35 | 0.833 |
| **Random Effects** | | | | | | | | |
| $\sigma^2$ | 3.29 | | | | 3.29 | | | |
| $\tau_{00}$ | 7.90 Household.file.id | | | | 11.95 Household.file.id | | | |
| | 1.14 sub_village.rnd | | | | 1.72 sub_village.rnd | | | |
| | 1.77 sub_village | | | | 2.27 sub_village | | | |
| | 0.59 ward.rnd | | | | 0.80 ward.rnd | | | |
| | 0.65 ward | | | | 0.00 ward | | | |
| | 0.22 district.rnd | | | | 0.00 district.rnd | | | |
| | 0.00 district | | | | 0.00 district | | | |
| N | 18067 Household.file.id | | | | 18067 Household.file.id | | | |
| | 2473 sub_village.rnd | | | | 2473 sub_village.rnd | | | |
| | 687 sub_village | | | | 687 sub_village | | | |
| | 663 ward.rnd | | | | 663 ward.rnd | | | |
| | 112 ward | | | | 112 ward | | | |
| | 54 district.rnd | | | | 54 district.rnd | | | |
| | 9 district | | | | 9 district | | | |
| Observations | 25677 | | | | 25677 | | | |
| Marginal $R^2$ / Conditional $R^2$ | 0.017 / 0.792 | | | | 0.060 / 0.846 | | | |
